# Aiptasia larvae are phenotypically validated as a model of coral bleaching using high-throughput machine-learning image analysis

**DOI:** 10.64898/2026.08.28.747729

**Authors:** Iyla Rossi, Emily K. Meier, Dania Nanes Sarfati, Francisco Guadalupe Zamora, Samantha Fung, Phillip A. Cleves, Amy E. Herr

## Abstract

The sea anemone Aiptasia is a model system for understanding cnidarian loss of symbiotic algae under heat stress (bleaching). While Aiptasia polyps have been widely used to study this process, accurate symbiosis phenotyping grapples with discordant length scales: fine spatial resolution (∼100 um) is needed across a whole organism (∼5 mm). To address this, we consider small (∼100 um), optically transparent Aiptasia larvae as a bleaching model suitable for whole-organism phenotyping by fluorescence microscopy with larvae classified as symbiotic when algae are localized within gastrodermal cells. To expedite phenotyping, we introduce a machine-learning (ML) image-analysis pipeline (SYMPHONY) designed for single-larva resolution analysis of intact larvae. SYMPHONY efficiently identifies the cellular location of internalized algae (accuracy: 79%, precision: 82%, recall: 79%, F1 score: 79%; training dataset composed of 1611 total objects). Additionally, SYMPHONY reports statistically significant larval bleaching under heat stress and corroborates manual phenotyping results, while significantly reducing operator labor from hours to minutes. The combination of the Aiptasia larvae model and the SYMPHONY pipeline aims to accelerate our understanding of symbiosis breakdown.

## Introduction

Coral reefs are foundational marine ecosystems, supporting exceptional biodiversity, protecting coastlines, and sustaining fisheries and tourism economies worldwide^1^. However, global climate change — particularly ocean warming — is driving the widespread collapse of coral reef ecosystems^2–4^. Specifically, coral bleaching occurs when environmental stress disrupts the mutualistic relationship between corals and their intracellular algae (family Symbiodiniaceae), leading to the loss of algae and their photosynthetically derived nutrients^5–7^. To understand how bleaching occurs, the sea anemone *Exaiptasia diaphana* (Aiptasia) has been utilized as a model system to study symbiosis formation and breakdown (bleaching)^8–10^. Aiptasia are related to reef-building corals and symbiose with the same strains of algae as corals but lack a calcium carbonate skeleton^5^. Additionally, Aiptasia are genetically tractable, grow rapidly, spawn frequently, and have a closed life cycle^8–12^. For these reasons, Aiptasia has become an important model system to study the molecular basis of cnidarian-algal symbiosis and its collapse^5,7^.

In particular, study of Aiptasia polyps (adults) has revealed genes associated with bleaching^7,13^. Nevertheless, methods to perturb (i.e., CRISPR/Cas9) and study gene function in a polyp can be time consuming, requiring nearly a year of husbandry to generate homozygous knockout polyps^14^. In addition, quantifying symbiosis in polyps can involve destructive sampling — lysis followed by normalization of algal cell counts to total host protein — or advanced imaging techniques (i.e., confocal microscopy, tissue sectioning) to assess physiology within tissue more than 100 µm thick^7,14,15^. While the former sacrifices spatial resolution and cannot distinguish between algae residing in symbiotic (gastroderm) versus non-symbiotic (gastric cavity) compartments, the latter is not readily scalable to phenotyping of an entire, intact anemone.

Consequently, Aiptasia larvae offer a promising complement to polyps as a model of coral bleaching. Larvae are small (∼100 µm in length and 50 µm wide), optically clear, and can be tested for gene function^5,14,16,17^. Owing to the transparent body of the larva, whole-larva imaging allows algal localization within the organism and quantitative assessment symbiosis phenotypes^16,17^. Previous studies report Aiptasia larvae as a suitable model to study the establishment of cnidarian-algal symbiosis^5^. However, an open question remains regarding whether Aiptasia larvae can serve as a model for bleaching, the breakdown of symbiosis^17^. Here, we show that, like polyps, Aiptasia larvae bleach and are suitable as a model for studying heat-stress induced bleaching. In tandem, we show that an ML pipeline incorporating single-cell mammalian (vs. cnidarian) biology tools performs image classification of confocal two-channel fluorescence images of larval morphology (staining, GFP spectral channel) and sub-larval algal localization (chlorophyll autofluorescence, Cy5 spectral channel). The “SYMPHONY” tool flattens the 3D fluorescence micrographs while retaining algae localization. Importantly, SYMPHONY standardizes micrograph preprocessing and performs 85% faster than human-expert image classification, with 79% accuracy. This study highlights Aiptasia larvae as a microscopy-compatible, efficient, and relevant model to study coral bleaching.

## Results

### Validating Aiptasia larvae as a bleaching model

Aiptasia larvae have a simple body plan with an outer epidermal tissue layer and inner gastrodermal layer that surrounds a gut-like cavity (gastric cavity). To establish symbiosis, algae must first enter the Aiptasia mouth and flow into the gastric cavity^5,18–21^. Once in the gastric cavity, Aiptasia gastrodermal cells can phagocytose algae into organelles that mature into a phagolysosome-like identity (symbiosome), wherein nutrient and metabolic exchange occurs^18–21^. This detail is important to how symbiosis is defined (**Fig. 1a**): algae are in symbiosis if they are within Aiptasia gastrodermal cells^14,21^. If algae are in the gastric cavity but not in the gastroderm, the Aiptasia is considered aposymbiotic. Cellular uptake of algae can occur at any time throughout the Aiptasia lifecycle, and the Aiptasia can contain algae in both the gastroderm and gastric cavity concurrently (in which case, the organism is symbiotic due to algae in the gastroderm)^14,21^. Therefore, localization of algae within the whole organism is necessary to determine the phenotype^14^.

**Fig. 1.**
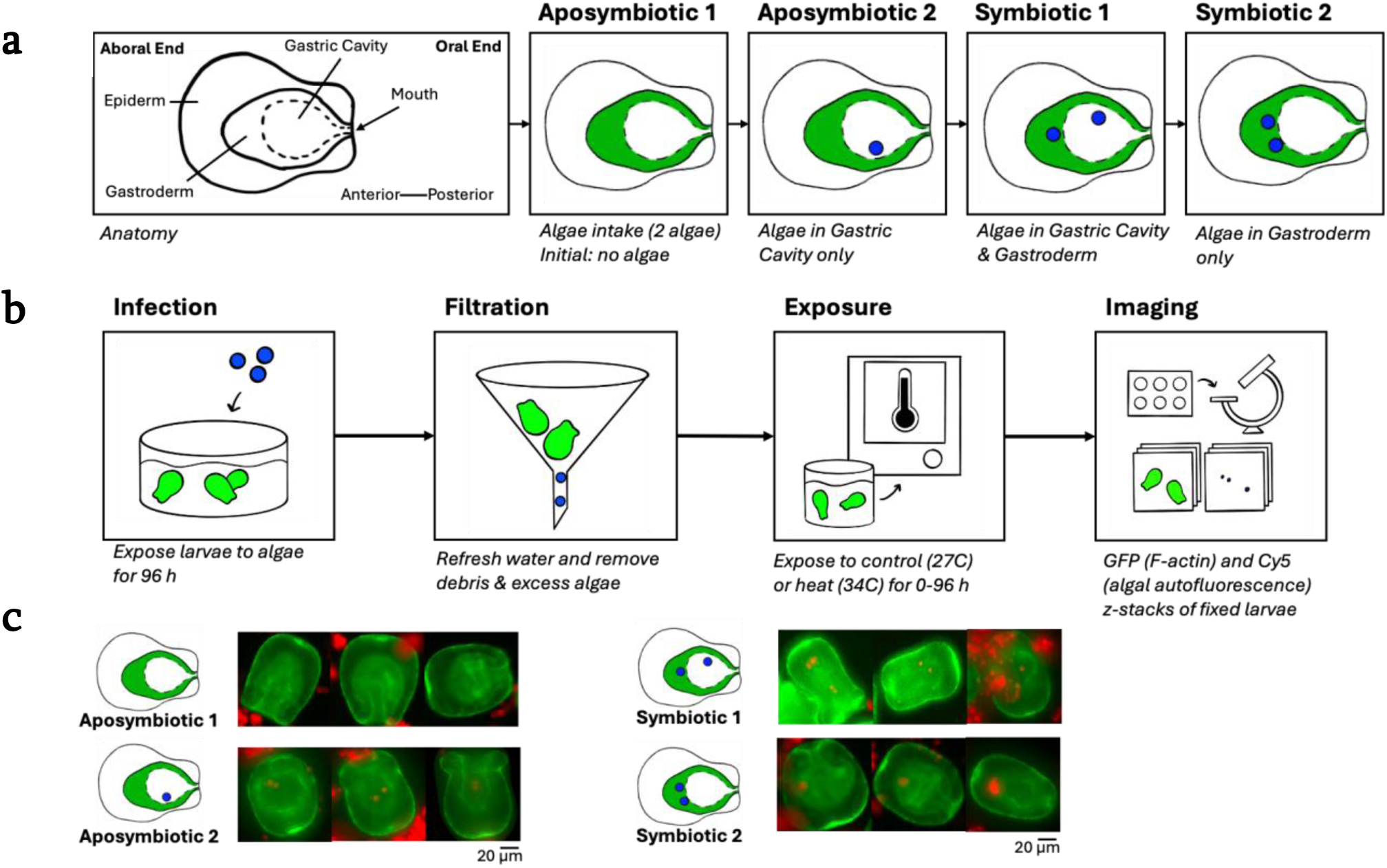
Anatomy, symbiotic phenotypes, and experimental workflow for Aiptasia larvae. **A** Diagram of larval anatomy showing the epidermis, gastroderm, and gastric cavity, along with anterior–posterior axes. The gastroderm surrounds the gastric cavity and forms the symbiotic tissue layer. **B** Diagrammatic overview of the experimental workflow. Larvae are exposed to Symbiodiniaceae algae (strain SSB01) for four days, filtered to remove algae and debris, subjected to algae-free temperature treatments, and imaged as dual-channel fluorescence z-stacks. **C** Representative fluorescence images corresponding to the four larval phenotypes included in the classification task. Aposymbiotic 1 larvae lack internal algae; Aposymbiotic 2 larvae contain algae localized to the gastric cavity; Symbiotic 1 larvae contain algae in both the gastric cavity and gastroderm; Symbiotic 2 larvae contain algae localized only within the gastroderm. Each row shows example images illustrating typical morphological and fluorescence features for that phenotype.

To investigate stress-induced bleaching in Aiptasia larvae, we performed a heat-stress assay (**Fig. 1b**), in a manner similar to a study that established Aiptasia polyps as a bleaching model^7^. We evaluated bleaching using three independent fertilizations to generate larvae “batches”, which were then infected with 150,000 cells/mL^-1^ of *Breviolum minutum* algae strain SSB01. Next, we exposed larvae to either a control (27 °C) or thermal stress (34 °C) condition for 24, 48, 72, or 96 h, with 0 h as the control^7^ and collected biological triplicates at each time point. We manually phenotyped 6,879 larvae within 81 fluorescence micrographs, requiring over 20.5 h of analysis. Representative images of four phenotypic classes — two aposymbiotic and two symbiotic — demonstrate the visual variability and biological distinctions at play (**Fig. 1c**). To understand bleaching rates in unpaired samples (i.e., each timepoint is an endpoint), we evaluated the change in the proportion of symbiotic larvae over the heat-stress time course. To approach this, we first enumerated on a per-well basis the number of symbiotic larvae (*N*_sym_) and aposymbiotic larvae (*N*_apo_) to obtain the proportion of symbiotic larvae (*P*_sym_) (equation 1):

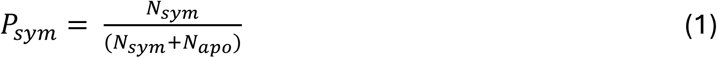

To normalize the proportion of symbiotic larvae from unpaired samples with multiple technical replicates (wells) per condition, we computed the weighted proportion of symbiotic larvae in the [0 h, 27℃] condition (C_sym_) for each experimental replicate:

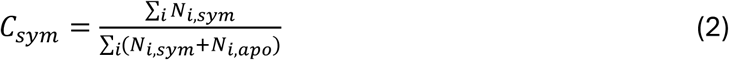

For each well, we next normalized *P*_sym_ to the *C*_sym_ from the same experimental replicate to obtain the proportion of remaining symbiotic larvae relative to the control proportion at [0 h, 27C], denoted as *P’*_sym_:

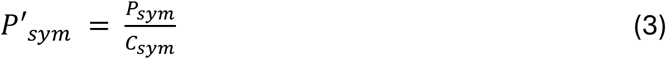

**Fig. 2** shows the normalized proportion of symbiotic larvae as a function of the duration of the temperature condition the larvae experienced.

**Fig. 2.**
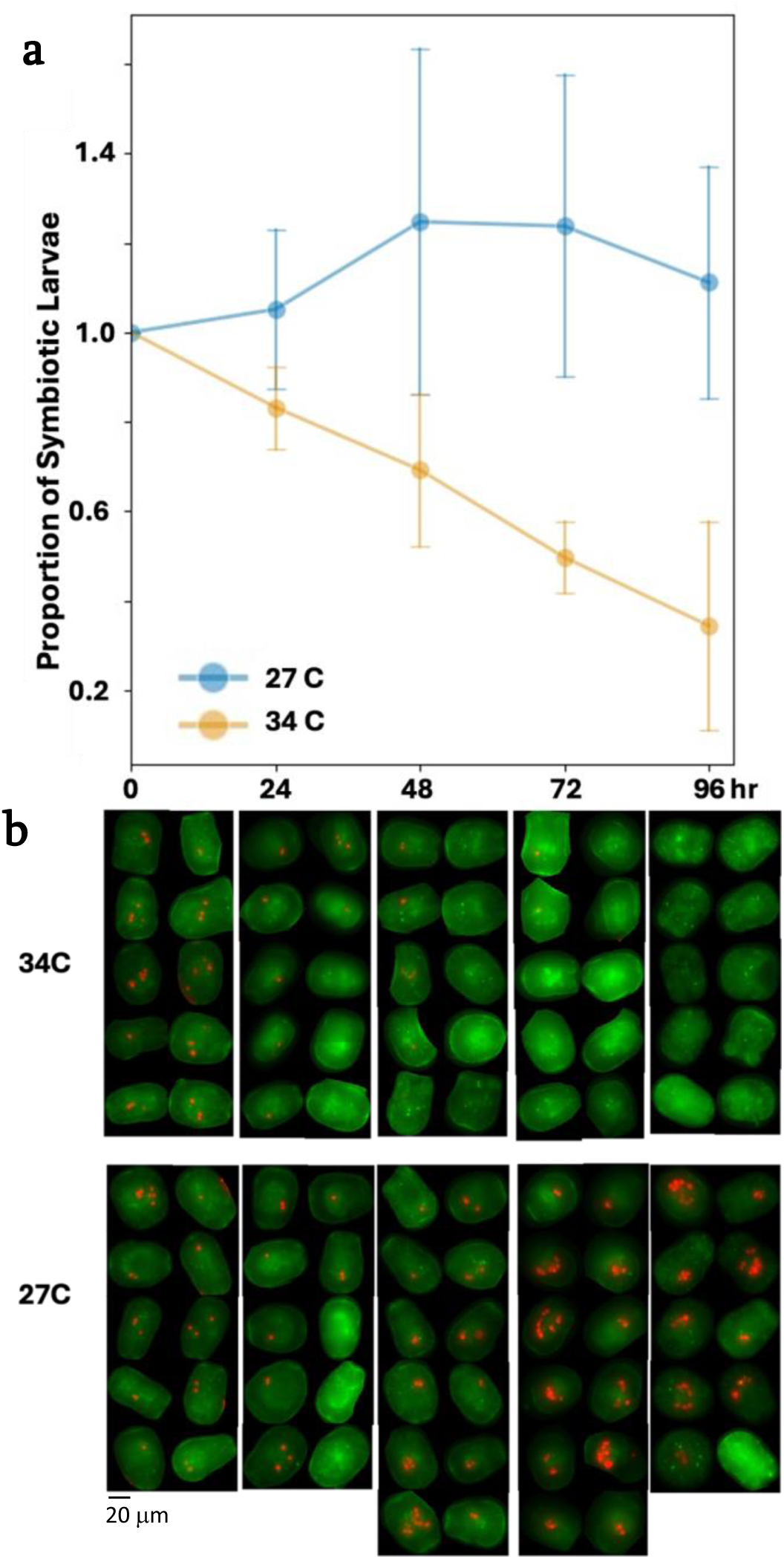
Symbiotic-state measurements and representative larvae across temperature and timepoints. **A** Proportion of larvae in symbiotic states at each temperature and timepoint (duration of exposure to the thermal condition). Data are normalized against the proportion of larvae in symbiotic states after 0 hrs of thermal exposure. **B** Representative cropped fluorescence images of larvae corresponding to each condition and timepoint, arranged so that their relative frequencies match the proportions observed in (a).

To test bleaching responses across timepoints accounting for batch effects in an unpaired longitudinal design, we used generalized estimating equation (GEE) regression with batch as a within-subject grouping factor. GEE provides a robust framework for analyzing proportion data derived from clustered observations. Unlike ANOVA, GEE does not require normally distributed response variables or balanced datasets. By specifying fertilization batch as a clustering variable, GEE enables population-averaged inference on the effects of temperature and duration on phenotype while remaining robust to potential misspecification of the within-batch correlation structure. By appropriately accounting for batch effects from fertilization, GEE allows us to leverage the full dataset across fertilization batches and experimental conditions to assess bleaching dynamics. The regression produced equation 4:

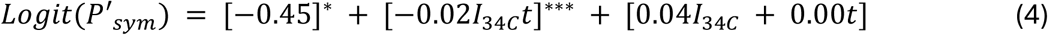

where *t* represents the duration of the thermal condition in hours and *I*_34C_ represents the intercept associated with the 34 °C condition, regardless of heat stress duration. Superscripts indicate the p-values associated with the coefficient (*** for p < 0.001; ** for p < 0.01; * for p < 0.05). The GEE regression indicates that, while there may be batch effects due to the experimental replicate, there is also a statistically significant decline in logit(P’_sym_) with increasing duration in heat stress (34 °C), but not with increasing duration in the 27 °C control condition. These results indicate that total algal loss (bleaching) increases with increasing duration in heat stress (34 °C). More specifically, we next asked whether the larvae follow the bleaching trends of previously studied adult Aiptasia bleaching systems. We observed Aiptasia larvae bleach ∼30% within 48 h (n=730, [-17%, 49%]) and ∼75% within 96 h (n=646, [24%, 81%]). Our observations show comparable results to a bleaching study performed on CC7 Aiptasia polyps with SSB01 algae following a similar heat exposure (34 °C heat condition, 27 °C control) which found ∼50% algal loss at 48 h and ∼100% algal loss at 96 h^10^. Showing similar findings, we moved forward in developing a pipeline to expedite the phenotyping process.

### Automating symbiosis phenotyping in Aiptasia larvae with the development of SYMPHONY

To streamline larvae symbiosis phenotyping, we developed an automated phenotyping pipeline to call symbiotic versus aposymbiotic larvae. This pipeline is applicable to z-stacks of 2-channel fluorescence images (i.e., the GFP channel for stained larvae tissue, and the Cy5 channel for algal chlorophyll autofluorescence). The pipeline was trained on a dataset consisting of 1,410 individual larvae. To build an automated pipeline, we first needed to preprocess micrographs containing multiple larvae.

First, multiple larvae in different symbiotic states (e.g., aposymbiotic, symbiotic) can be observed within the same fluorescence micrograph. Image classification is often the preferred approach and is widely implemented in microscopy workflows; however, it can only assign a single label to the whole image and lacks region- or pixel-level granularity. In our case, because we have multiple larvae in different symbiotic states per micrograph, we opted to implement instance segmentation to identify the pixel-level locations and individual phenotypes of the larvae^22–26^. Since manual annotation requirements are time-consuming for instance segmentation, we sought to minimize this burden by layering image classification (You Only Look Once, YOLO^27^) on top of an existing generalist segmentation algorithm (Cellpose^28^).

Second, larvae anatomy is three dimensional. Consequently, the micrographs must also be three-dimensional and are therefore acquired as epi-fluorescence microscopy z-stacks. 3D object detection and instance segmentation typically require annotating/drawing the locations of each larva in each image across the entire 3D image z-stack^28^. This 3D processing notably increases the time needed for data annotation^28^. Further, as the model complexity increases with 3D data, 3D data typically requires larger training datasets than 2D data^28^. To minimize data complexity and annotation requirements, we first reduce data dimensionality without losing axial information by projecting a 3D image into a 2D representation. Third, we aimed to minimize training needs and model complexity by leveraging preexisting ML models.

We designed and developed a custom pipeline, SYMPHONY (<u>Sy</u>mbiosis <u>M</u>achine-learning-based <u>PH</u>en<u>O</u>typing and a<u>N</u>al<u>Y</u>sis), containing three modules to address the complexities of larvae symbiosis phenotyping (**Fig. 3**). To reduce the dimensionality of our dataset while preserving axial (3D) information, our first module encodes axial information into a compact 2D representation. Next, to minimize data annotation requirements, our second module leverages Cellpose, a segmentation algorithm from single-cell mammalian biology, to identify and isolate individual larvae into their own images^28^. Finally, to determine the larval phenotypes, our third module leverages YOLO, a preexisting image classification architecture, to assign and count phenotypes from the larvae isolated in module 2 ^27^. Below, each module is described in detail.

**Fig. 3.**
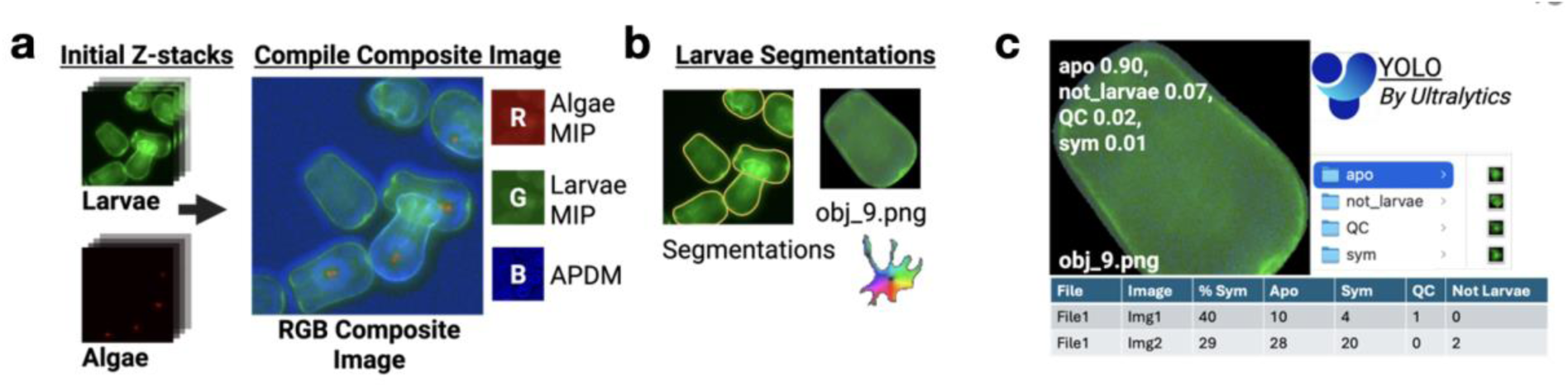
Overview of the SYMPHONY depth-aware phenotyping pipeline. **A** Preprocessing workflow illustrating the three images derived from each z-stack: the F-actin maximum-intensity projection (G channel), the algal autofluorescence maximum-intensity projection (R channel), and the axial position difference map (B channel). These channels are merged to create a depth-aware RGB composite. **B** Example segmentation output showing a single larva isolated from the merged composite. A Cellpose-generated mask is overlaid to indicate the extracted region of interest used for downstream cropping. **C** Classification interface and output. The extracted larval crop is shown alongside the predicted probabilities of the classifier for each phenotype category. The bottom panel shows aggregated classification summaries for two example image files, including counts and percentages of larvae assigned to each class.

### Module 1: Depth-aware preprocessing preserves axial information lost in 2D projections

To minimize the training dataset needs and computational complexity of the pipeline, we computed maximum intensity projections (MIPs). These MIPs store only the maximum intensity (i.e., the brightest pixel) from the entire z-stack for each (x, y) coordinate^29^. We computed MIPs for both the larvae GFP and algae Cy5 channels, allowing us to reduce the dimensionality from 3D to 2D. However, MIPs obscure axial context by collapsing fluorescence from multiple z-planes into a single 2D image^29^. As shown in **Fig. 4a**, a larva containing algae in both the gastroderm and gastric cavity (biologically symbiotic) can appear identical to a larva containing algae only in the gastric cavity (biologically aposymbiotic) when viewed as a 2D projection. This visual ambiguity represents a fundamental limitation of MIP and other projection-based approaches for developing computational tools to phenotype larvae symbiosis.

**Fig. 4.**
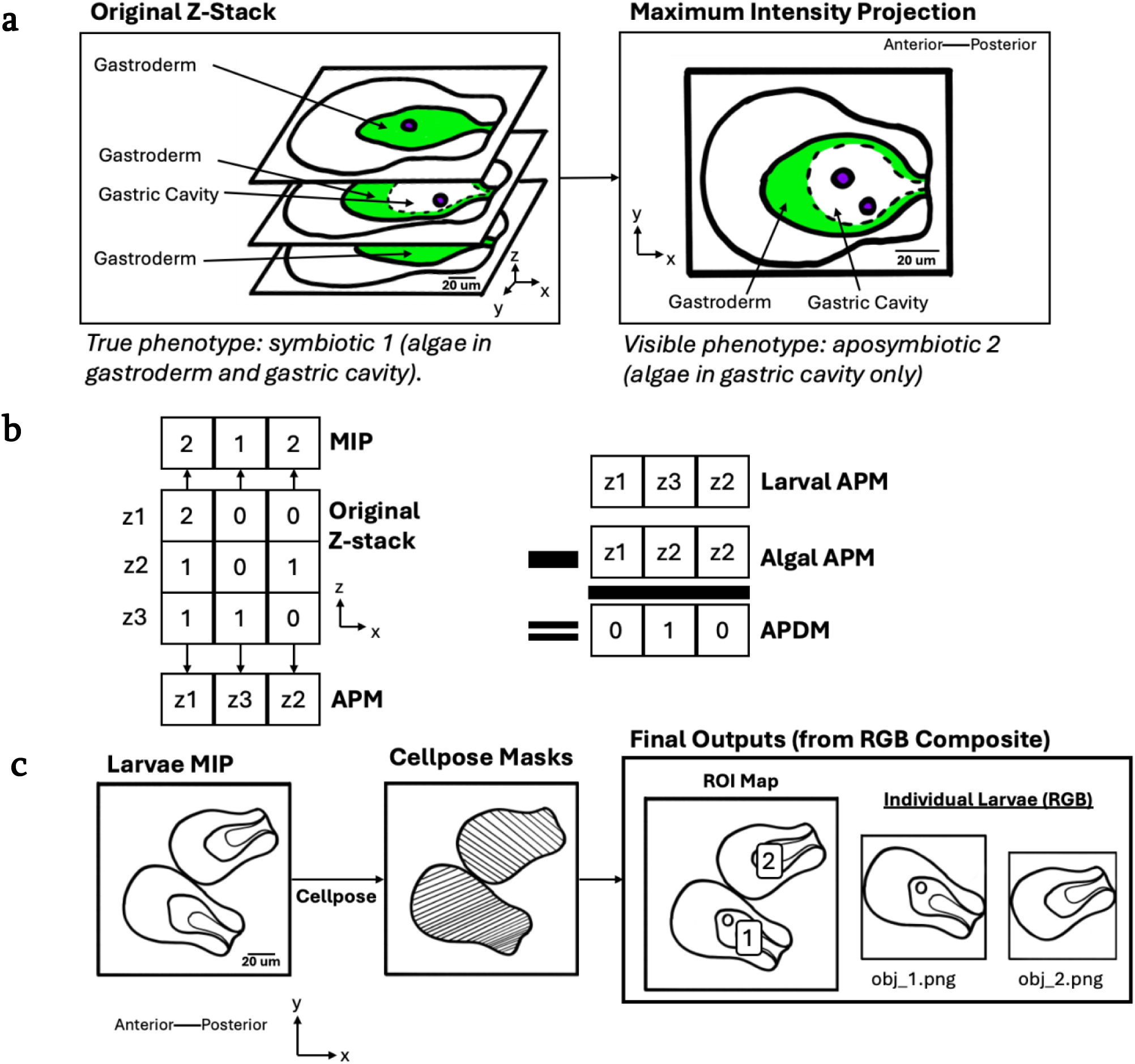
Depth-aware representation and segmentation steps used to prepare larvae for classification. **A** Schematic illustrating the effect of maximum-intensity projection on axial information. A multipanel stack of larval z-slices is shown alongside the resulting 2D projection. **B** Illustration of how pixel-wise axial information is derived. Z-slices are arranged in a matrix, and the z-position of maximal intensity is determined for each pixel to generate axial position maps and axial position difference maps. **C** Steps used to segment and crop individual larvae. Left: Maximum-intensity composite images of multiple larvae. Middle: Corresponding segmentation masks produced by Cellpose. Right: Region-of-interest maps and final cropped RGB images used as classifier inputs.

To preserve axial structure, SYMPHONY uniquely determines the argmax – the z-position from which the maximum intensity pixel was derived – for every (x, y) coordinate. The full array of these z-positions is henceforth referred to as the axial position map (APM). By computing the MIPs and APMs, the original 3D fluorescence z-stack images were compressed to a set of four 2D images, each corresponding to a single fluorescence channel (i.e., MIP_algal_, MIP_larval_, APM_algal_, and APM_larval_).

Next, we aimed to use preexisting image segmentation and classification algorithms to identify and phenotype larvae from the projection images. However, most existing image-based algorithms are designed for RGB images, which encode colors as the combination of additive primaries, storing three intensity values per (x, y) coordinate: [red intensity, green intensity, blue intensity]. Since each of our 2D fluorescence projection images (MIPs and APMs) correspond to a single fluorescence channel, each projection image is grayscale, containing only one intensity value per pixel instead of the three intensity values found in RGB images. However, by combining the four projections, we have four intensity values per (x, y) coordinate instead of the three-value format allowed by RGB-based algorithms. To ensure compatibility for downstream use in module 3 (which uses preexisting RGB-based image classification algorithms requiring three intensity values per coordinate), we aimed to match the dimensionality of our 2D fluorescence projection images to a standard 2D RGB image.

One way to address this is by computing the axial distance between the algae and the larval gastroderm to phenotype symbiosis, as the positions of the organisms relative to the image or microscope are not necessarily needed. To compute the axial distance between algae and larvae for every (x, y) coordinate, SYMPHONY subtracts the algal APM from the larval APM. This mapping, henceforth called the axial position difference map (APDM), resolves the issue of our previous four intensity projection map as we now have three: MIP_algal_, MIP_larval_, and APDM. Now with three projections, SYMPHONY generates a standard 2D RGB image by encoding each projection as a single color. The red intensities encode algal data and correspond to the MIP of the Cy5 channel (MIP_algal_). The green intensities, encoding larvae data, correspond to the MIP of the GFP channel (MIP_larval_). Finally, the blue intensities, encoding the distance between algae and larvae, correspond to the APDM. **Fig. 4b** demonstrates how the original fluorescence data maps to this final 2D RGB image, henceforth called the RGB composite image. The RGB composite image reduces the dimensionality of our 3D dataset, simplifies data complexity and annotation requirements, and preserves awareness of the axial positioning of algae relative to the larvae gastroderm for each xy coordinate. Further, this RGB composite image is compatible with existing algorithms for image-based ML (e.g., commercial YOLO models).

Axial information is crucial for phenotyping larvae-algal symbiosis. Therefore, we explored using existing depth-based RGB ML algorithms for rapid phenotyping. Depth encodings have previously been employed for rapid data visualization and ML in biological and biomedical imaging^30^. Although such color-coded MIPs efficiently communicate depth information for single-channel datasets (e.g., fluorescence or brightfield), they are not well suited to multi-channel fluorescence imaging. MIPs have also been combined with depth encodings to generate 2D renderings of volumetric data using depth-based shading, in which intensity values are modulated to produce shadowing effects that enhance 3D perception^31^. However, these shading-based approaches may compromise interpretability for quantitative phenotyping, as fluorescence intensity (encoded in the MIP) is itself biologically meaningful. More recently, projection-based intensity representations have been paired with depth encodings in optical coherence tomography^32^. While this work establishes precedent for integrating projection-based intensity maps with depth information, it fundamentally differs from fluorescence microscopy in both signal origin and contrast mechanism^33^. These approaches remain limited in the context of multi-channel fluorescence imaging, where simultaneous preservation of channel identity and axial relationships is required, as in our need, for proper phenotyping.

Using the APDM, SYMPHONY extends previous depth-encoded projection strategies to encode the relative axial displacement between two biologically distinct and meaningful fluorescence channels, rather than absolute depth with respect to the imaging axis. This relative-depth representation improves generalizability and model accuracy across micrographs acquired regardless of changes in the number of or distance between focal planes. In addition, individual larvae may occupy distinct axial planes within a single field of view, further underscoring the limitations of absolute depth encoding and motivating a relative approach.

By fusing channel-specific MIPs with a biologically meaningful relative-depth encoding into a standard 2D RGB representation, SYMPHONY preserves the spatial relationships that define symbiosis phenotypes while remaining compact, interpretable, and compatible with existing 2D ML algorithms. This framework establishes a generalizable design strategy for integrating multi-channel volumetric biological data into conventional 2D learning pipelines without sacrificing biological interpretability.

### Module 2: Generalist segmentation achieves robust larval isolation

As described previously, our original micrographs and RGB composite images may contain multiple larvae per image, requiring object detection or instance segmentation. To balance the need for detection or segmentation with the small size of our training dataset (1,410 total larvae: 722 aposymbiotic, 474 symbiotic, 214 quality control or borderline phenotypes) and the desire to minimize manual annotation, we employed Cellpose, a generalist algorithm trained to segment individual mammalian cells, in our non-cell based context^28,34^. As a generalist algorithm, Cellpose has been trained on thousands of images of various cell types, primarily but not limited to mammalian cell types, improving its generalizability^28,34^. Additionally, Cellpose resizes images based on the expected pixel diameter of the object of interest, and not by physical length dimensions of the cells^28^. Together, these factors have enabled Cellpose as a tool applied in non-cell-based contexts, including segmenting worms and autophagic bodies in yeast vacuoles^28,34–36^. Further, Cellpose has been trained to distinguish nuclei from cytoplasms in both brightfield and fluorescence micrographs^28^. Together, because our fluorescent micrographs show clearly defined discoid shapes, we can implement Cellpose to treat larvae as ‘cytoplasms’ and segment out single larvae from an image that may contain multiple.

We employ the Cellpose ‘cytoplasm’ model on the larval MIP (GFP) to identify the pixels corresponding to each individual larva, regardless of phenotype. **Fig. 4c** depicts this process. This allows us to minimize our annotation and training dataset requirements, as de-novo construction and training of a model to segment the larvae is not needed. Instead, we can leverage the Cellpose segmentations to form new images containing only one larva each, thereby converting a segmentation problem into a classification problem to simplify our training data requirements.

To create images containing only one larva each, we create a copy of the RGB composite image for each larva identified by Cellpose. Then, for the n-th identified larva, we modify the n-th copy of the RGB composite image, named ‘obj_n.png’ for traceability. To isolate the n-th larva, we set all pixels not corresponding to the n-th larva (these may correspond to background or other larvae) to black. To reduce the size of the image and the overall dataset, we crop the image to the nearest square that encompasses and centers the n-th larva. The square crop also ensures that images can be rescaled as needed for downstream classification algorithms without altering the larva. By centering the cropped images on the larvae, we minimize positional bias; we ensure the cropped square is centered on the larva by padding the image with extra black pixels as needed. These final images are henceforth referred to as individual larval images (ILIs). For traceability, we also generate an ROI map, in which the ‘cell’ number is labeled on the original RGB composite image so the user can locate the origin of each ILI.

Thus, our second module leverages Cellpose for multicellular marine biology, converting a custom segmentation task into a traceable 2-step process to balance segmentation requirements with small datasets and annotation availability: (1) phenotype-unaware general segmentation and (2) classification.

### Module 3: Classification distinguishes larvae symbiosis phenotypes

For phenotype classification, we fine-tuned the commercial Ultralytics YOLO11 model on SYMPHONY’s individual larval images (ILIs)^27^. Ultralytics created the first Python-based YOLO model, and Ultralytics YOLO models have been previously used for cell detection and tracking in microscopy^37–40^.

We trained this classifier to assign each ILI to one of four categories: aposymbiotic, symbiotic, quality control (QC), and not_larva. **Fig. 5a** shows representative examples of each category in which the number of examples shown reflects the prevalence of that phenotype in the available training dataset. The QC class captures malformed larvae, atypical orientations (i.e., larvae facing up or down), or cases where fluorescence signal is insufficient for reliable assignment. Finally, the not_larva class captures any non-larval objects identified by Cellpose, such as bubbles, debris, or image artifacts.

**Fig. 5.**
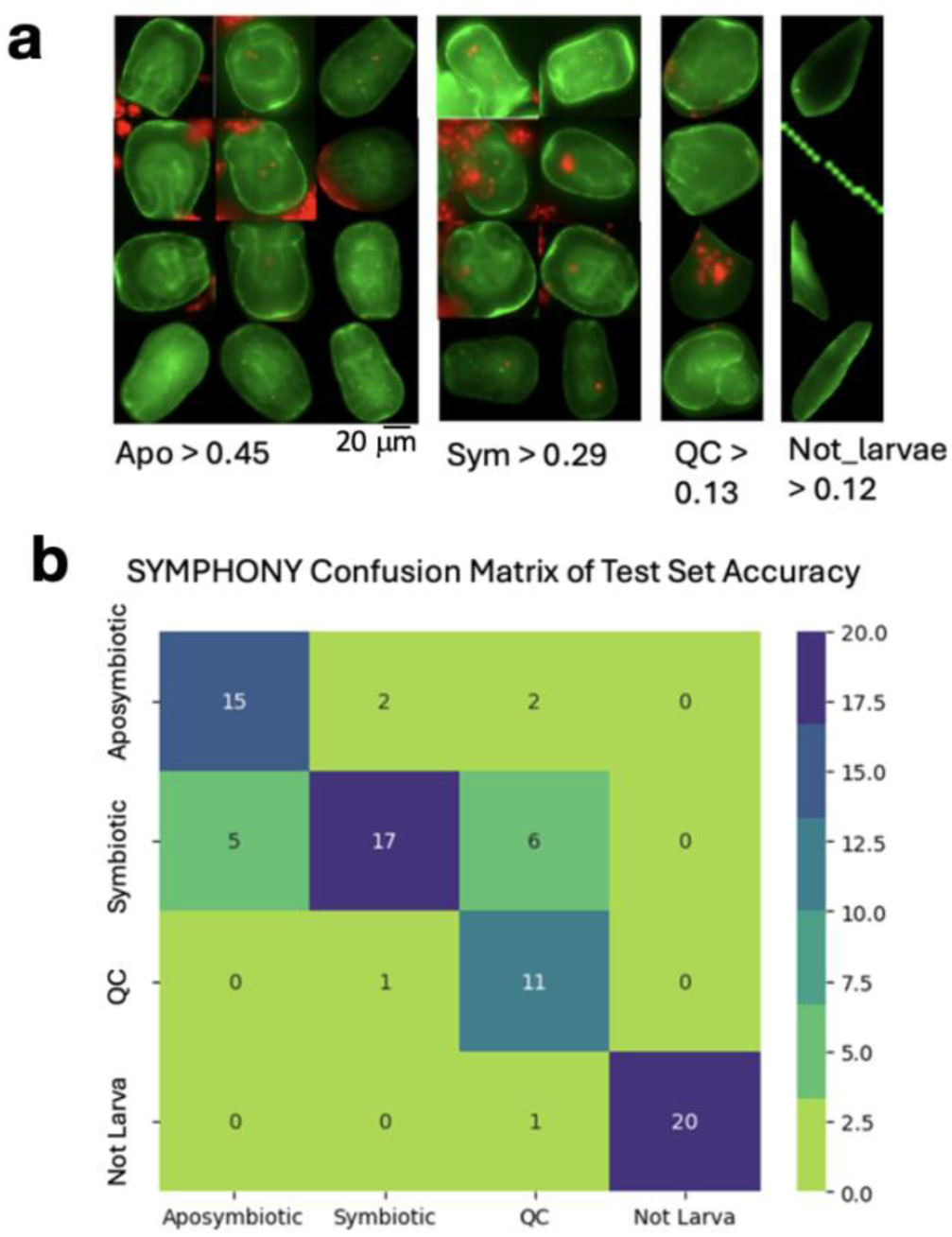
Representative classification inputs and confusion matrix for SYMPHONY’s four-class model. **A** Representative cropped images from each of the four phenotype classes. Each panel shows multiple larvae displaying the range of intensities, shapes, and fluorescence patterns present within the training set. **B** Confusion matrix summarizing classifier output for the four phenotype categories. Rows represent true labels and columns represent predicted labels. Color values denote the number of images in each classification outcome.

Following classification, we sort the ILIs into folders based on predicted phenotype. We next count the number of ILIs per phenotype to obtain the number of larvae per phenotype from the original 3D image.

Our dataset consisted of 1,611 ILIs: 722 aposymbiotic, 474 symbiotic, 214 QC, and 201 not_larva. **Fig. 5a** shows representative images from each class in the dataset. Because our phenotypes (classes) vary in the number of ILIs, we have an imbalanced dataset, which can lead to model predictions based on class sizes rather than image content^41–44^. For example, as our dataset contains significantly fewer symbiotic ILIs than aposymbiotic ILIs, the model might learn to never predict a larva has the symbiotic phenotype. Additionally, small datasets risk overfitting, in which the model learns predictions based on the specific noise in the dataset, reducing its applicability to new data^45^.

To mitigate class imbalance and overfitting, we used Albumentations, a Python package for custom image augmentations, which has been used for medical diagnostic models and microscopy-based ML^46–52^. To prevent overfitting, we split our dataset into training, test, and validation sets before augmentation. Specifically, we extracted 20 images per phenotype into each of the test and validation sets to enforce a class balance in test and validation metrics.

To form the training dataset, we performed image augmentation on the remaining images until all classes contained an equal number of images (598 training images per phenotype, 2,392 total training images). Additionally, we designed our augmentation pipeline with biological constraints in mind. First, we cannot accurately phenotype a larva with no algae visible in the gastroderm when significant portions of the gastroderm are missing from the image, as these regions may contain symbiotic algae. Thus, such larvae should be classified as ‘QC’. However, if a symbiotic ILI undergoes spatial transformation during augmentation such that the regions of the gastroderm containing algae are no longer present, the resulting ILI would be misclassified as ‘symbiotic’ when it should be classified as ‘QC’. To prevent such phenotype alterations and misclassifications caused by augmentation, spatial transformations were restricted to minimize significant loss of larval data.

Furthermore, due to the relationship between the MIPs (red=algae chlorophyll autofluorescence in Cy5; green=larvae F-actin stain in GFP) and the APDM (blue channel, larvae depth – algae depth), we imposed constraints on the augmentations to preserve biological feasibility and relevance. Through augmentation we oversample underrepresented classes by duplicating images from those classes to increase the number of images per class^53^. Further, the augmentation process introduced variation in the dataset by altering these duplicates to improve generalizability.

To evaluate classifier performance, we computed a confusion matrix summarizing prediction accuracy across all classes (**Fig. 5b**). The classifier achieved an overall accuracy of 79% and high recall for both aposymbiotic and symbiotic classes, with most misclassifications occurring between ambiguous larvae and the QC category. This pattern is expected given that QC images represent borderline cases by design. To further characterize classifier performance, we computed per-class accuracy. We also computed the precision, or the proportion of positive predictions that were correct^54^. Additionally, we computed the recall, which represents the model’s ability to correctly identify positive cases of a class^54^. Finally, we computed the F1 score, the harmonic mean of precision and recall^54^. All metrics are reported in **Table 1**. The model achieved high recall for symbiotic (0.85) and aposymbiotic (0.75) larvae, with strong precision across all classes (macro precision = 0.82). QC larvae displayed lower recall (0.55) but high precision (0.92), reflecting the conservative behavior of the classifier when encountering borderline or low-signal images. Not_larva predictions were uniformly accurate (accuracy = 1.0; recall = 1.0), indicating reliable removal of non-biological inputs from downstream phenotyping analysis.

**Table 1.** SYMPHONY performance metrics across each phenotype.

| Class | Accuracy | Precision | Recall | F1 |
| --- | --- | --- | --- | --- |
| Aposymbiotic | 0.75 | 0.79 | 0.75 | 0.77 |
| Symbiotic | 0.85 | 0.61 | 0.85 | 0.71 |
| QC | 0.55 | 0.92 | 0.55 | 0.69 |
| Not_larva | 1 | 0.95 | 1 | 0.98 |
| Macro | 0.79 | 0.82 | 0.79 | 0.79 |

### SYMPHONY reports larvae bleach under heat stress

To test the ability of the SYMPHONY automated micrograph analysis pipeline to efficiently and effectively phenotype symbiosis, we assessed SYMPHONY’s ability to characterize larvae bleaching during persistent heat stress. We again computed the proportion of symbiotic larvae per well and the normalized proportion relative to the (27 °C x 0 h) condition. **Fig. 6** shows the SYMPHONY-computed results. Similarly, we performed a GEE regression on the results computed by SYMPHONY:

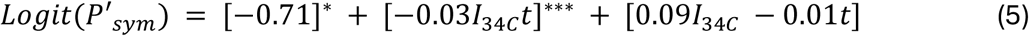

where *t* represents the duration of the thermal condition and *I*_34C_ represents the intercept associated with the 34 °C condition, regardless of heat stress duration. The superscripts indicate the p-values associated with the coefficient: *** p < 0.001, ** p < 0.01, * p < 0.05. The decrease in logit(P’_sym_) due to continued exposure of 34 °C is statistically significant (p < 0.001, n = 89, clusters = 3), while the change due to temperature or time alone are not (temperature: p = 0.474; time: p = 0.069; n = 89, clusters = 3). SYMPHONY indicates that larval bleaching occurs due to continued exposure to heat stress (34 °C).

**Fig. 6.**
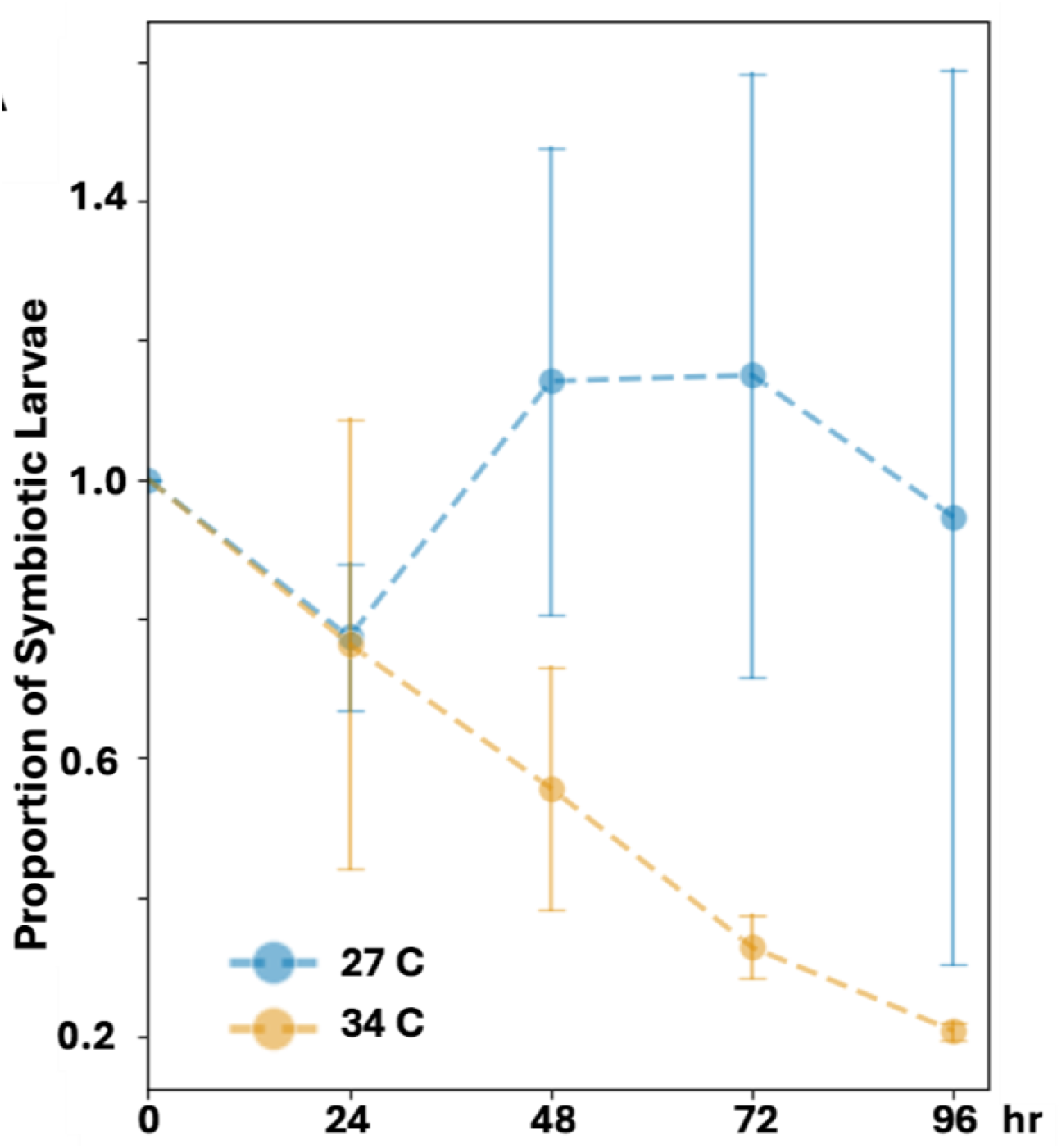
Automated symbiotic-state measurements. Proportion of larvae classified into symbiotic states at each temperature and timepoint (duration of exposure to the thermal condition). Data are normalized against the proportion of larvae in symbiotic states after 0 h of thermal exposure.

Next, we compared SYMPHONY’s results to our previous manual results validating larvae as a bleaching model (**Fig. 7**). We see that the coefficient on the bleaching term (I_34C_t) is consistent across the human- and SYMPHONY-based analyses, and nearly identical (-0.02 and -0.03, respectively).

**Fig. 7.**
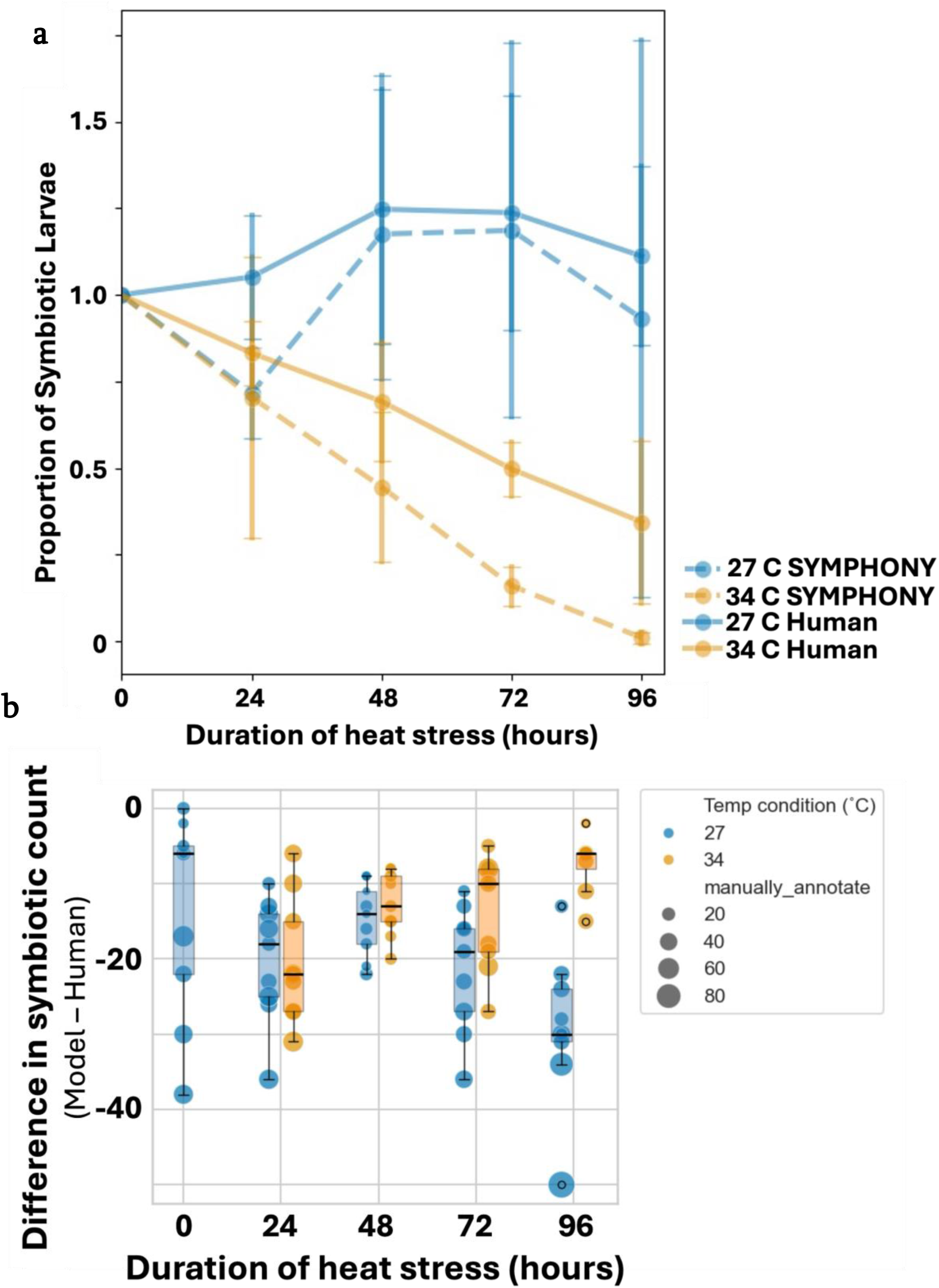
Comparison of symbiotic-state estimates and relationship between QC assignments and model deviations. **A** Symbiotic-state proportions plotted across temperature and timepoint conditions. Data are shown for both human annotations and automated classifier outputs. **B** Scatter and boxplot depiction of the difference between classifier and human symbiotic counts. Point size is proportional to the number of larvae labeled as ‘QC’ within each sample.

To further evaluate SYMPHONY’s ability to recapitulate the manual results, we performed another GEE regression on the data, this time including method (SYMPHONY vs. human) as a term:

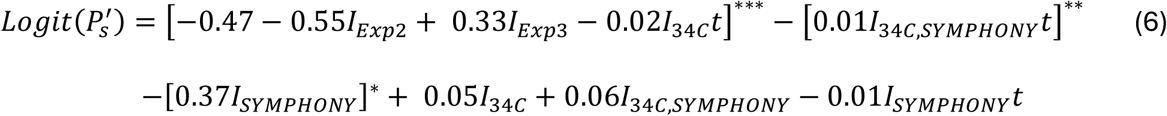

where *t* represents the duration of the thermal condition and *I*_34C_ represents the intercept associated with the 34 °C condition, regardless of heat stress duration. The superscripts indicate the p-values associated with the coefficient: *** p < 0.001, ** p < 0.01, * p < 0.05.

We observed statistically significant bleaching due to the term −0.02*I_34C_t* (p<0.001, n=178, clusters=89). However, the results also indicate that SYMPHONY over-reports bleaching (−0.01*I_34C_*_,*SYMPHONY*_*t*) and underreports symbiosis (-0.37*I_SYMPHONY_*). We attribute this phenomenon to the QC phenotype, which flags larvae with difficult-to-label phenotypes due to orientation or position in the z-stack, as larvae classified as QC are flagged for human analysis and excluded from the computation of P_sym_. To investigate SYMPHONY’s underreporting of symbiosis, we graphed the deviation between the number of symbiotic larvae reported by SYMPHONY and human analysis (**Fig. 7**). The results indicate that images with severe underreporting of symbiosis contained a significant number of larvae classified as QC, supporting the finding that SYMPHONY recapitulates the results from our previous validation of larval bleaching. Based on prior literature, we hypothesize that training on a dataset containing at least 150 examples per phenotypic sub-class (e.g., symbiotic larvae with different orientations or algal localization patterns) would reduce the number of phenotypes classified as QC, thereby mitigating the underreporting of symbiosis^55^. With greater use/data input, SYMPHONY could be retrained to learn to phenotype some QC larvae.

Finally, we compared the computational time of SYMPHONY to the time required for manual annotation validation of larval bleaching. SYMPHONY requires ∼ 30 s to set up and begin phenotyping for a dataset. After the initial 30 s of operator “manual” labor, SYMPHONY required 3.2 h to complete the automated phenotyping (of 355.95 GB of input data). In contrast, manual phenotyping required over 20 h of labor. If images flagged for QC by SYMPHONY are not manually phenotyped, SYMPHONY saved over 20 h of manual labor for our study, or ∼ 9.33 s / larva. However, if QC images are later phenotyped manually, with 11 s required to phenotype each larva, SYMPHONY saves over 12 h of manual labor, or 7 s / larva.

## Discussion

This study consisted of two major advancements, the first assessing the suitability of Aiptasia larvae as a bleaching model, and the second developing new tools for symbiosis research. To understand the suitability of Aiptasia larvae as a model of coral-algal bleaching, we evaluated larval bleaching under heat stress. Adult Aiptasia polyps have previously been used as a bleaching model and have provided insights into genomic and transcriptomic changes during coral bleaching^7,56^. We validated Aiptasia larvae as a bleaching model and implemented the SYMPHONY pipeline to address the challenges of phenotyping symbiosis in a rapid, high-throughput manner.

The first advancement aimed to use a simplified system to study symbiosis and bleaching, and one compatible with scalable and high-throughput experimentation. We turned to the Aiptasia larvae system due to their small size and optical clarity, weekly access to thousands of larvae from polyp spawning, and the ability of the larvae to rapidly take up algae. Together, these features made the larvae system attractive to develop high-throughput phenotyping methods.

Next, we aimed to validate Aiptasia larvae as a bleaching model. We asked if Aiptasia larvae bleach in a similar manner to polyps. Manual phenotyping of heat-stressed larvae found a significant, 40% decrease in symbiosis at 48 hr in heat stress and 75% decrease in symbiosis after 96 hr in 34 °C heat stress compared to the 27 °C control. Though the trajectory of algal loss in larvae is not as dramatic as that seen in polyps harboring the same algal species (reporting nearly 100% loss of algae in 96 hr), the bleaching trend is largely similar and notably distinct from bleaching trends observed with polyps harboring other algal species^7,56–60^. Because Aiptasia larvae and polyps exhibit similar loss of SSB01 algae, the larvae represent a viable model to study bleaching. Similarities between larvae and polyp heat-induced bleaching should be investigated and may reveal core, conserved molecular programs involved in heat stress response and algae loss. By establishing Aiptasia larvae as a bleaching model, microscopy-based phenotyping and quantitative analysis of intact whole Aiptasia is possible. Further, Aiptasia larvae are suitable for scalable and high-throughput experiments in combination with SYMPHONY.

In our second advancement, we developed SYMPHONY, an automated pipeline to phenotype Aiptasia larvae through a three-module process leveraging pre-existing tools from mammalian biology: (1) data collapse from 3D to 2D, (2) larval segmentation and isolation into individual images, (3) phenotype classification and results aggregation. We designed and deployed SYMPHONY with a graphical user interface (GUI) that opens in a default browser window initiated by a single line of code, regardless of operating system. Additionally, closing the GUI window automatically shuts down all code and processing, minimizing the amount of code or programming experience required to operate the GUI. The GUI contains options to access files from your local desktop, specify a GPU or CPU, and modify select parameters. Finally, the GUI and code are open access via a GitHub repository that includes download instructions for installation and usage.

Using our SYMPHONY pipeline, symbiosis phenotyping from an entire experiment can occur in just 10 days, from the time of fertilization to imaging the experiment. Further, a wide variety of genetic methods are already established for Aiptasia larvae^18,61^. These methods include the use of short hairpin RNA knockdown, CRISPR/Cas9 mutagenesis, and plasmid DNA and mRNA overexpression^61,62^. These methods could be employed in our pipeline to test Aiptasia genes that function in bleaching^7^. Furthermore, the pipeline could be adapted as a high-throughput method to study Aiptasia genes that affect the formation and maintenance of the symbiosis^61^. Lastly, SYMPHONY could be used to assess differences in bleaching rates in Aiptasia larvae harboring different algal symbiont genotypes. Thus, by establishing Aiptasia larvae as a bleaching model with rapid phenotyping, we aim to accelerate studies examining the mechanisms underlying heat-induced bleaching.

Looking ahead, SYMPHONY is designed to be scalable for image-based phenotyping in high-throughput experiments (e.g., gene knockout screens, large-scale environmental perturbation studies, and the development of diagnostic assays to assess bleaching susceptibility and resilience). We opted for an open-source framework and modular design to provide a template for extending automated phenotyping to other symbiotic systems, accelerating discovery at the intersection of developmental biology, ecology, and environmental genomics.

With this current assay and SYMPHONY pipeline, we assess changes in larval symbiosis under persistent heat stress. Because this sampling uses fixed larvae sampled from each timepoint, we are assessing the effect of heat stress on a population of larvae (proportion symbiotic statistic). This is useful for accounting for variability of symbiosis effects within a population, but lacks the ability to track and observe the effect of heat stress on single larvae over time. New imaging techniques, such as high resolution particle tracking and live imaging, could enable future work that builds on the SYMPHONY pipeline to observe and characterize symbiosis breakdown from persistent heat stress in single larvae over time. This might yield greater information about the mechanisms by which heat-induced bleaching occurs. Furthermore, because SYMPHONY can be retrained and greatly reduces operator labor, SYMPHONY could be adapted to look at other dynamics of the symbiosis, including but not limited to: formation and maintenance of the symbiosis, perturbation of these states, or differences in these processes owed to different Aiptasia or algal genotypes. In each case, high-throughput analyses may contribute to a greater understanding of what parameters contribute to supporting the symbiosis, with implications for coral conservation.

## Materials and Methods

### Heat Stress Experiments

#### Aiptasia Animal Husbandry and Generation of Wildtype Aiptasia larvae

Adult Aiptasia polyp colonies CC7 and PLF3 clonal populations infected with SSB01 algae were reared in individual 250 mL tanks with ∼33 ppt artificial seawater (ASW) (Saltwater Aquarium’s Coral Pro Salt Mix – Red Sea) in Percival incubators at 27 °C with 25 µmol photons m^-2^ s^-1^ full spectrum white light under a 12:12h light:dark schedule. Polyps were fed *Artemia* brine shrimp twice a week and water changed on the day of feeding. Cohorts of CC7 and PLF3 polyps were induced to spawn in flipped incubators with a moonlight cue using previously described methods^63^. Four cohorts offset by one week allowed for weekly spawning (given the monthly gametogenic cycle). CC7 sperm from the 250 mL tanks was filtered to remove tank debris using a 75 µm filter and collected in a 1 L glass beaker. PLF3 eggs were collected around the base around PLF3 polyps using disposable 2 mL transfer pipets and transferred into a 250 mL straight sided, flat bottom glass bowl containing ∼5 mL ASW. Once all eggs were collected, ∼200 mL of filtered CC7 sperm was added to the glass bowl containing eggs to generate wildtype Aiptasia larvae. A clear, round plastic cover was used to cover the bowl and prevent evaporation throughout the duration of the experiments. We then placed the covered bowl into the 27 °C incubator to allow for zygote development. 24 h later, larvae were filtered using a 40 µm mesh filter and washed gently yet thoroughly with a squeeze-bottle containing 0.22 µm sterile filtered ASW (FASW) to remove debris and dead cells. The cleaned larvae were then moved into a fresh glass bowl with 200 mL FASW, and placed back into a 27 °C Percival incubator with a plastic cover.

### Methods of Culturing Breviolum minitum Algae Strain SSB01

Algae of *Breviolum minitum* strain SSB01 were grown in Daigo’s IMK medium (ASW with 252 mg L^−1^ Daigo’s IMK powder) (FujiFilm barcode no. 4987481531246). The IMK medium was sterilized using a 0.1-µm filter and grown at the same temperature and light Percival incubator settings as Aiptasia, described previously. Algae were passaged every month in a 1:10 dilution using 200 mL tissue culture flasks with vent caps (Cell Treat, Cat. No. 229341).

### Aiptasia Larvae Infections with Breviolum minitum Strain SSB01

Algae cells from the flask were pelleted at 1 min 3,000 x g in 50 mL Falcon tubes, washed thrice with FASW, and quantified using a Luminex Guava EasyCyte flow cytometer (Cytek Biosciences).

2 days post fertilization (dpf), Aiptasia larvae were filtered using a 40 µm mesh filter, thoroughly rinsed with FASW, and concentrated into 5 mL FASW in preparation for infection. The larvae concentration was checked by averaging triplicate larvae counts in 100 µL aliquots. Larvae were evenly split across 3 straight sided, flat bottom glass bowls (∼2,000 larvae bowl^-1^) and inoculated with 250 mL of FASW containing 150,000 SSB01 algae mL^-1^. The glass bowls containing larvae and algae were covered with a clear plastic cover to prevent evaporation, and the larvae were allowed to infect for 4 days under control temperature (27 °C).

### Division into Heat Stress Assay Conditions

At 6 dpf, the ASW containing larvae and algae was passed through two stacked mesh filters, a 150 µm and 40 µm mesh, to remove algae clumps objects larger than the larvae and to capture the larvae, respectively. Larvae captured on the 40 µm mesh filter were thoroughly rinsed with FASW to remove excess algae and debris. Finally, larvae were divided equally into glass bowls containing 200 mL FASW for either control (27 °C) or heat stress (34 °C) treatments, and covered with a clear piece of plastic to prevent evaporation.

We selected 27 °C and 34 °C as our control and thermal stress conditions, respectively, in line with a previous Aiptasia bleaching study^7^. Our temperatures fall within the validated ranges for Aiptasia husbandry (25-27 °C) and thermal stress (31-34 °C)^7,64–68^. Temperature and light were monitored via HOBO pendant loggers (Onset Computer Corporation Cat.

No. UA-002-64). Salinity was stable at 32-33 ppt throughout the duration of the experiments in both heat stress and control conditions, as measured by a MA887 digital seawater refractometer (Milwaukee, Cat. No. MA887) before and after the heat stress experiment.

Larvae were exposed to their given condition for 24, 48, 72, or 96 h, with 0 h of exposure serving as the control^7^. At each heat-stress timepoint, 100 larvae per condition were sampled in triplicate (total 300 larvae per condition timepoint).

### Heat Stress Assay Replicates

For robustness, we performed the complete bleaching assay on larvae from three separate spawning fertilization dates (3-26-2025, 4-02-2025, 7-09-2025), yielding three experimental replicates, hereafter referred to as ‘batches.’ Within each batch, larvae were divided into 3 glass bowls for each (temperature x duration) condition, with ∼ 100 larvae sampled per dish. Thus, across 3 batches, there were 9 samples per (temperature × duration) condition, subject to batch effects. Additionally, the division of larvae into wells occurred before the bleaching time course, so the larvae in (27 °C x 24 h) were independent of those in (27 °C x 96 h) within a given batch.

### Fixation and Imaging Protocol

At each heat-stress timepoint, three replicates of 100 larvae per temperature condition were sampled in 40 µL aliquots 4-well dishes (Cell Treat Cat. No. 229503) and chemically fixed with 40 µL of glutaraldehyde and paraformaldehyde solution (0.1% and 4% final concentrations, respectively) in FASW for 5 min at room temperature. The first fix mixture was removed and larvae were re-fixed with 80 µL of 4% PFA in FASW for 1 h at 4 °C. Larvae were then washed 5× with cold 1× PBSTw (1x PBS with 0.1% Tween-20) in 5 minute incubations on a rotator, before staining for 1 h with Phalloidin-488 (final concentration 1:3000 from 400x stock) and Hoechst 33342-405 (final concentration 1 µg/mL) in 1× PBS at 4 °C, out of light. Samples were washed 2 times with cold 1x PBSTw for 15 min at room temperature on a rotator before mounting. SecureSeal imaging spacers (8-9 mm diameter, 0.12 mm depth) were used for mounting larvae, with each replicate placed in an individual well. Larvae in 1x PBSTw were moved into a well, the 1x PBSTw media removed, and mounted in 4.5 µL of 80% glycerol/1x PBS and covered with a #1.5 cover slip.

A Leica DM6 M upright widefield epi-fluorescence microscope with an automatic stage imaged the larvae using the automatic stage set for the SecureSeal imaging spacers. Images were taken using the 20x objective, 30% FIM, and tiled z-stacks centered around the mean depth with the focus map calibrated to the middle z-stack of each individual well. Tiled z-stack images were collected in both GFP and Cy5 spectral channels and stitched to create one xyz image per well. The Cy5 fluorescence was imaged at 125 ms exposure, and the GFP was imaged at 700 ms exposure. All experiments were imaged using the same exposure and z-stack settings.

### SYMPHONY Pipeline

#### Software and Hardware

All analysis was performed on a 2022 Apple MacBook Air (M2, 16GB RAM, macOS Sonoma 14.1.2). The pipeline used Python 3.11 with PyTorch, Ultralytics, Cellpose, Panel, and scientific computing libraries. Full version details and dependencies are provided below (*Methods: SYMPHONY Pipeline: Software Package Versions*). ChatGPT was used to assist with debugging code.

#### SYMPHONY Pipeline Overview

Z-stacks (GFP and Cy5) were processed into RGB images encoding larval and algal maximum intensity projections (MIPs; green, red) and the axial position difference map (blue). Cellpose segmented larvae from the larval MIP, and bounding boxes and segmentation masks were used to crop individual larval images. YOLO classified these into phenotypes. Aggregated outputs include phenotype counts and symbiotic proportions.

#### Training Dataset and Augmentations

Training data consisted of PLF3 eggs crossed to CC7 sperm to generate wildtype Aiptasia larvae that were infected at 2 dpf for 4 days with SSB01 algae and fixed 6 dpf. Larvae were fixed, stained, and imaged in a manner similar to that noted previously (*Methods: Heat Stress Assay: Fixation and Imaging Protocol*), with the exception that both 1:6000 and 1:3000 Phalloidin-488 stain were used to provide a measure of variability in the training data to ensure the robustness of the SYMPHONY image-analysis tool. The stitched and merged GFP and Cy5 z-stack training data images were processed to the cropping step. RGB crops were manually labeled in Piximi into four phenotype classes (apo, sym, manually_annotate, not_larvae). Labeled images were organized into train, test, and validation folders compatible with YOLO using a custom Python script, and the training dataset was augmented with Albumentations.

#### Module I: Image Preprocessing

Images were loaded via readlif (for LIF) or tifffile (for TIFF). MIPs were generated using NumPy. Axial Position Maps (APMs) captured z-indices of intensity maxima. Axial Position Difference Maps (APDMs) were computed as larval APM − algal APM. RGB composites used R = algae MIP, G = larval MIP, B = APDM.

#### Module II: Larval Segmentation

Cellpose v3.1 with ‘cyto3’ model was used on larval MIP. A fixed object diameter of 140 µm was converted to pixel units. Segmentation masks were exported and used to crop larvae with bounding box logic ensuring square aspect ratio. Pixels outside the mask were zeroed to prevent background interference.

Bounding boxes were determined for each segmentation mask. To enforce square crops, the shorter side (width or height) was symmetrically extended to match the longer side. If the resulting box extended beyond image bounds, the crop was padded with black ([0, 0, 0]) pixels. Only pixels within the mask preserved their original RGB values; background pixels were zeroed. Each larva crop was saved in TIFF and PNG formats with filenames preserving traceability to the source image and mask ID.

#### Module III: Phenotype Classification

Cropped larvae were classified with YOLOv11 (Ultralytics v8.0.2) into four classes (apo, sym, QC, not_larvae). Output images were saved with the classes ranked by confidence scores superimposed upon the image. These output images were saved in folders based on the Top-1 predicted class.

To aggregate these results, we create a spreadsheet with columns for the filename, original image name, phenotypes, and summary statistics, such as the proportion of symbiotic larvae. If the spreadsheet already exists (e.g., when the pipeline is applied to multiple files at once), the data is appended as a new row. Otherwise, a new spreadsheet is created.

#### GUI Functionality and Deployment

The GUI was developed using Panel and launched with a single terminal command. It includes four tabs: Home, Output Folders, Editable Parameters, and Logs. GUI logs pipeline activity and enables batch processing.

### Software Package Versions

numpy==1.24.4

pandas==2.1.3

scikit-image==0.25.1

opencv-python==4.10.0

tifffile==2023.7.10

zarr==2.15.0

matplotlib==3.10.0

PyYAML==6.0.1

tqdm==4.66.1

torch==2.5.1

ultralytics==8.0.200

cellpose==3.1.0

scikit-learn==1.6.1

readlif==0.6.2

xmltodict==0.14.2

panel==1.3.0

An environment.yml and requirements.txt are available on GitHub.

## Acknowledgements

We are grateful to the Symbiosis in Aquatic Systems Initiative from the Gordon and Betty Moore Foundation for their funding and support (awarded to P.A.C. and A.E.H.). Additionally, we thank the NIH training program T32GM139780 for their funding and support (awarded to I.R.).

## Author Contributions

I.R., E.K.M., D.N.S., P.A.C., and A.E.H. designed the study. E.K.M. performed the experiments and generated the epi-fluorescence micrographs. I.R. developed the SYMPHONY pipeline. I.R. generated the SYMPHONY training data from the epi-fluorescence micrographs. I.R., E.K.M., F.G.Z., and S.F. labeled the training data. I.R. and E.K.M. wrote the original draft of the manuscript. D.N.S., F.G.Z., S.F., P.A.C., and A.E.H. reviewed and edited the manuscript.

## References

1. Chen, P.-Y., Chen, C.-C., Chu, L. & McCarl, B. Evaluating the economic damage of climate change on global coral reefs. Global Environmental Change 30, 12–20 (2015).

2. Hughes, T. P. et al. Coral reefs in the Anthropocene. Nature 546, 82–90 (2017).

3. Hoegh-Guldberg, O. et al. Coral Reefs Under Rapid Climate Change and Ocean Acidification. Science 318, 1737–1742 (2007).

4. Jackson, J. B. C. Ecological extinction and evolution in the brave new ocean. Proceedings of the National Academy of Sciences 105, 11458–11465 (2008).

5. Wolfowicz, I. et al. Aiptasia sp. larvae as a model to reveal mechanisms of symbiont selection in cnidarians. Sci Rep 6, 32366 (2016).

6. Roth, M. S. The engine of the reef: photobiology of the coral–algal symbiosis. Front Microbiol 5, 422 (2014).

7. Cleves, P. A., Krediet, C. J., Lehnert, E. M., Onishi, M. & Pringle, J. R. Insights into coral bleaching under heat stress from analysis of gene expression in a sea anemone model system. Proceedings of the National Academy of Sciences 117, 28906–28917 (2020).

8. Weis, V. M., Davy, S. K., Hoegh-Guldberg, O., Rodriguez-Lanetty, M. & Pringle, J. R. Cell biology in model systems as the key to understanding corals. Trends in Ecology & Evolution 23, 369–376 (2008).

9. Baumgarten, S. et al. The genome of Aiptasia, a sea anemone model for coral symbiosis. Proc Natl Acad Sci U S A 112, 11893–11898 (2015).

10. Sunagawa, S. et al. Generation and analysis of transcriptomic resources for a model system on the rise: the sea anemone Aiptasia pallida and its dinoflagellate endosymbiont. BMC Genomics 10, 258 (2009).

11. Lehnert, E. M., Burriesci, M. S. & Pringle, J. R. Developing the anemone Aiptasia as a tractable model for cnidarian-dinoflagellate symbiosis: the transcriptome of aposymbiotic A. pallida. BMC Genomics 13, 271 (2012).

12. Grawunder, D. et al. Induction of Gametogenesis in the Cnidarian Endosymbiosis Model Aiptasia sp. Sci Rep 5, 15677 (2015).

13. Weizman, E. & Levy, O. The role of chromatin dynamics under global warming response in the symbiotic coral model Aiptasia. Commun Biol 2, 282 (2019).

14. Tran, C. et al. Photosynthesis and other factors affecting the establishment and maintenance of cnidarian–dinoflagellate symbiosis. Philosophical Transactions of the Royal Society B: Biological Sciences 379, 20230079 (2024).

15. Tivey, T. R., Coleman, T. J. & Weis, V. M. Spatial and Temporal Patterns of Symbiont Colonization and Loss During Bleaching in the Model Sea Anemone Aiptasia. Front. Mar. Sci. 9, (2022).

16. Hambleton, E. A., Guse, A. & Pringle, J. R. Similar specificities of symbiont uptake by adults and larvae in an anemone model system for coral biology. J Exp Biol 217, 1613–1619 (2014).

17. Van Treuren, W. et al. Live imaging of Aiptasia larvae, a model system for coral and anemone bleaching, using a simple microfluidic device. Sci Rep 9, 9275 (2019).

18. Bucher, M., Wolfowicz, I., Voss, P. A., Hambleton, E. A. & Guse, A. Development and Symbiosis Establishment in the Cnidarian Endosymbiosis Model Aiptasia sp. Sci Rep 6, 19867 (2016).

19. Xiang, T. et al. Symbiont population control by host-symbiont metabolic interaction in Symbiodiniaceae-cnidarian associations. Nat Commun 11, 108 (2020).

20. Rädecker, N. et al. Using Aiptasia as a Model to Study Metabolic Interactions in Cnidarian-Symbiodinium Symbioses. Front. Physiol. 9, (2018).

21. Jinkerson, R. E. et al. Cnidarian-Symbiodiniaceae symbiosis establishment is independent of photosynthesis. Current Biology 32, 2402–2415.e4 (2022).

22. Hafiz, A. M. & Bhat, G. M. A survey on instance segmentation: state of the art. Int J Multimed Info Retr 9, 171–189 (2020).

23. Su, H. et al. Object Detection and Instance Segmentation in Remote Sensing Imagery Based on Precise Mask R-CNN. in IGARSS 2019 - 2019 IEEE International Geoscience and Remote Sensing Symposium 1454–1457 (2019). doi:10.1109/IGARSS.2019.8898573.

24. Zou, Z., Chen, K., Shi, Z., Guo, Y. & Ye, J. Object Detection in 20 Years: A Survey. Proceedings of the IEEE 111, 257–276 (2023).

25. Druzhkov, P. N. & Kustikova, V. D. A survey of deep learning methods and software tools for image classification and object detection. Pattern Recognit. Image Anal. 26, 9–15 (2016).

26. M.V., A. & Khan, D. M. Recent Trends on Object Detection and Image Classification: A Review. in 2020 International Conference on Computational Performance Evaluation (ComPE) 427–435 (2020). doi:10.1109/ComPE49325.2020.9200080.

27. Ultralytics. Ultralytics YOLO11. https://docs.ultralytics.com/models/yolo11/.

28. Stringer, C., Wang, T., Michaelos, M. & Pachitariu, M. Cellpose: a generalist algorithm for cellular segmentation. Nat Methods 18, 100–106 (2021).

29. Fishman, E. K. et al. Volume Rendering versus Maximum Intensity Projection in CT Angiography: What Works Best, When, and Why. RadioGraphics 26, 905–922 (2006).

30. Otsuna, H., Ito, M. & Kawase, T. Color depth MIP mask search: a new tool to expedite Split-GAL4 creation. Preprint at 10.1101/318006 (2018).

31. Zhou, Z., Tao, Y., Lin, H., Dong, F. & Clapworthy, G. Shape-enhanced maximum intensity projection. Vis Comput 27, 677–686 (2011).

32. Thrapp, A. D. et al. Feasibility of Depth-in-Color En Face Optical Coherence Tomography for Colorectal Polyp Classification Using Ensemble Learning and Score-Level Fusion. J Biophotonics 19, e202500292 (2026).

33. Yang, C. Molecular Contrast Optical Coherence Tomography: A Review. Photochemistry and Photobiology 81, 215–237 (2005).

34. Stringer, C. & Pachitariu, M. Cellpose3: one-click image restoration for improved cellular segmentation. 2024.02.10.579780 Preprint at 10.1101/2024.02.10.579780 (2024).

35. Marron, E. C., Backues, J., Ross, A. M. & Backues, S. K. Accurate automated segmentation of autophagic bodies in yeast vacuoles using cellpose 2.0. Autophagy 20, 2092– 2099 (2024).

36. Wheeler, N. J., et al. wrmXpress: A modular package for high-throughput image analysis of parasitic and free-living worms. PLOS Neglected Tropical Diseases 16, e0010937 (2022).

37. Al-Hamadani, M. N. A. et al. Improving Cell Detection and Tracking in Microscopy Images Using YOLO and an Enhanced DeepSORT Algorithm. Sensors 25, (2025).

38. Mehta, P. et al. Benchmarking YOLO Variants for Enhanced Blood Cell Detection. International Journal of Imaging Systems and Technology 35, e70037 (2025).

39. Nabiullina, R. et al. 3D Bioprinting of Cultivated Meat Followed by the Development of a Fine-Tuned YOLO Model for the Detection and Counting of Lipoblasts, Fibroblasts, and Myogenic Cells. Front. Biosci. (Landmark Ed) 30, (2025).

40. Pascual-González, M., Jiménez-Partinen, A., Palomo, E. J., López-Rubio, E. & Ortega-Gómez, A. Hyperparameter optimization of YOLO models for invasive coronary angiography lesion detection and assessment. Computers in Biology and Medicine 196, 110697 (2025).

41. Chang, C.-Y., Hsu, M.-T., Esposito, E. X. & Tseng, Y. J. Oversampling to Overcome Overfitting: Exploring the Relationship between Data Set Composition, Molecular Descriptors, and Predictive Modeling Methods. J. Chem. Inf. Model. 53, 958–971 (2013).

42. Li, Z., Kamnitsas, K. & Glocker, B. Overfitting of Neural Nets Under Class Imbalance: Analysis and Improvements for Segmentation. in Medical Image Computing and Computer Assisted Intervention – MICCAI 2019 (eds. Shen, D. et al.) 402–410 (Springer International Publishing, Cham, 2019). doi:10.1007/978-3-030-32248-9_45.

43. Santos, M. S., Soares, J. P., Abreu, P. H., Araujo, H. & Santos, J. Cross-Validation for Imbalanced Datasets: Avoiding Overoptimistic and Overfitting Approaches [Research Frontier]. IEEE Computational Intelligence Magazine 13, 59–76 (2018).

44. Mujahid, M. et al. Data oversampling and imbalanced datasets: an investigation of performance for machine learning and feature engineering. J Big Data 11, 87 (2024).

45. Ying, X. An Overview of Overfitting and its Solutions. J. Phys.: Conf. Ser. 1168, 022022 (2019).

46. Buslaev, A. et al. Albumentations: Fast and Flexible Image Augmentations. Information 11, (2020).

47. Kamardi, C. et al. Classification of Alzheimer’s Disease using Random Oversampling and Albumentations on Convolutional Neural Network. in 2023 Eighth International Conference on Informatics and Computing (ICIC) 1–6 (2023). doi:10.1109/ICIC60109.2023.10382106.

48. Lee, M.-H. et al. Alignment-free bacterial pathogen identification using vision transformer and image augmentation techniques in high-resolution microscopy. Biomedical Signal Processing and Control 112, 108401 (2026).

49. Sifat, M. M. A. et al. A ResNet50 Transfer Learning and Grad-CAM-Based Framework for Explainable TEM Nanoparticle Classification. Preprint at 10.21203/rs.3.rs-7933991/v1 (2025).

50. Marques, G., Ferreras, A. & de la Torre-Diez, I. An ensemble-based approach for automated medical diagnosis of malaria using EfficientNet. Multimed Tools Appl 81, 28061– 28078 (2022).

51. Hallström, E., Kandavalli, V., Wählby, C. & Hast, A. Rapid label-free identification of seven bacterial species using microfluidics, single-cell time-lapse phase-contrast microscopy, and deep learning-based image and video classification. PLOS ONE 20, e0330265 (2025).

52. Papa, M., Bhattacharya, S., Park, B. & Yi, J. Rapid Salmonella Serovar Classification Using AI-Enabled Hyperspectral Microscopy with Enhanced Data Preprocessing and Multimodal Fusion. Foods 14, (2025).

53. Shorten, C. & Khoshgoftaar, T. M. A survey on Image Data Augmentation for Deep Learning. J Big Data 6, 60 (2019).

54. Sujon, K. M., Hassan, R., Choi, K. & Samad, M. A. Accuracy, precision, recall, f1-score, or MCC? empirical evidence from advanced statistics, ML, and XAI for evaluating business predictive models. J Big Data 12, 268 (2025).

55. Shahinfar, S., Meek, P. & Falzon, G. “How many images do I need?” Understanding how sample size per class affects deep learning model performance metrics for balanced designs in autonomous wildlife monitoring. Ecological Informatics 57, 101085 (2020).

56. Natural Variation in Responses to Acute Heat and Cold Stress in a Sea Anemone Model System for Coral Bleaching | The Biological Bulletin: Vol 233, No 2. *The Biological Bulletin* https://www.journals.uchicago.edu/doi/10.1086/694890.

57. Gegner, H. M. et al. High levels of floridoside at high salinity link osmoadaptation with bleaching susceptibility in the cnidarian-algal endosymbiosis. Biol Open 8, bio045591 (2019).

58. Sydnor, J. R., Lopez, J., Wolfe, G. V., Ott, L. & Tran, C. Changes in the microbiome of the sea anemone Exaiptasia diaphana during bleaching from short-term thermal elevation. Front. Mar. Sci. 10, (2023).

59. Gegner, H. M. et al. High salinity conveys thermotolerance in the coral model Aiptasia. Biol Open 6, 1943–1948 (2017).

60. Bieri, T., Onishi, M., Xiang, T., Grossman, A. R. & Pringle, J. R. Relative Contributions of Various Cellular Mechanisms to Loss of Algae during Cnidarian Bleaching. PLOS ONE 11, e0152693 (2016).

61. Renicke, C. et al. Development of genetic tools for the sea anemone Aiptasia, a model system for coral biology. Genetics 231, iyaf194 (2025).

62. Maruyama, S. et al. Co-option of lysosomal machinery shapes the symbiosis supporting coral reefs. 2025.10.09.679812 Preprint at 10.1101/2025.10.09.679812 (2025).

63. Lab, P., Perez, S. & Barry, O. Aiptasia spawning and embryo/larvae handling - Pringle Lab. https://www.protocols.io/view/aiptasia-spawning-and-embryo-larvae-handling-pring-x54v98edml3e/v1 (2018).

64. Allen-Waller, L. R., Jones, K. G., Martynek, M. P., Brown, K. T. & Barott, K. L. Comparative physiology reveals heat stress disrupts acid–base homeostasis independent of symbiotic state in the model cnidarian Exaiptasia diaphana. J Exp Biol 227, jeb246222 (2024).

65. Ishii, Y. et al. Global Shifts in Gene Expression Profiles Accompanied with Environmental Changes in Cnidarian-Dinoflagellate Endosymbiosis. G3 (Bethesda) 9, 2337–2347 (2019).

66. Randle, J. L., Cárdenas, A., Gegner, H. M., Ziegler, M. & Voolstra, C. R. Salinity-Conveyed Thermotolerance in the Coral Model Aiptasia Is Accompanied by Distinct Changes of the Bacterial Microbiome. Front. Mar. Sci. 7, (2020).

67. Herrera, M. et al. Temperature transcends partner specificity in the symbiosis establishment of a cnidarian. ISME J 15, 141–153 (2021).

68. Mansfield, K. M. et al. Transcription factor NF-κB is modulated by symbiotic status in a sea anemone model of cnidarian bleaching. Sci Rep 7, 16025 (2017).

